# vCA1 SST neurons represent avoidance states that guide anxiety-related behavioral choices

**DOI:** 10.64898/2026.03.02.709156

**Authors:** Joshua X Bratsch-Prince, Mingyang Wei, Srutika Sureshbabu, Vivan Chhaya, Nuria Cano Adamuz, Kelsey Logas, Mazen A. Kheirbek

**Affiliations:** Department of Psychiatry and Behavioral Sciences, University of California, San Francisco, San Francisco, CA, USA; Neuroscience Graduate Program, University of California, San Francisco, San Francisco, CA, USA; Kavli Institute for Fundamental Neuroscience, Weill Institute for Neurosciences and Center for Integrative Neuroscience, University of California, San Francisco, San Francisco, CA, USA

## Abstract

In uncertain, threatening environments, rapid and flexible behavioral decisions are essential for survival. The ventral CA1 region of the hippocampus encodes threatening contexts and can regulate avoidance-related behaviors. While activity patterns in excitatory neurons of vCA1 have been well characterized, the contributions of specific interneuron subtypes to avoidance decisions in threatening situations remain to be fully elucidated. Here, we show that somatostatin expressing (SST) interneurons in vCA1 are selectively recruited during avoidance behavior and play a causal role in shaping behavioral responses in aversive spaces. We find that vCA1 SST interneurons ramp their activity in anticipation of, and during, avoidance. This contrasted with parvalbumin (PV) and vasoactive intestinal peptide (VIP) interneurons, which are preferentially active during exploratory approach behaviors. Unlike other classes of vCA1 interneurons, SST neuron activity more reliably represented the animal’s intention to avoid rather than its spatial position. Moreover, optogenetic silencing of SST interneurons reduced the efficiency of approach-avoidance decisions. These findings identify vCA1 SST interneurons as key regulators of threat assessment, revealing a cell-type-specific mechanism by which vCA1 microcircuits govern avoidance behaviors. This work provides a new framework for understanding hippocampal control of avoidance behavior and highlights SST-expressing interneurons as key contributors to anxiety-related behaviors.

## Introduction

The ventral hippocampus, particularly ventral CA1 (vCA1), is a critical node in an extended circuit that processes salient emotional and contextual stimuli (Biane et al., 2025; Fanselow and Dong, 2010; Kjelstrup et al., 2002; Turner et al., 2022; Xia et al., 2025). While there is extensive evidence implicating vCA1 pyramidal cells (PCs) in representing anxiety-provoking environments and modulating behavioral responses to anxiogenic environments (Adhikari et al., 2011, 2010; Ciocchi et al., 2015; Jimenez et al., 2020, 2018; Padilla-Coreano et al., 2019; Parfitt et al., 2017; Sánchez-Bellot and MacAskill, 2020; Trent and Menard, 2010; Yeates et al., 2022), how discrete circuit elements within this structure represent distinct features of anxiety-like states, such as features of the environment, internal emotional states, or behavioral states of avoidance, remain poorly understood.

The vCA1 microcircuit contains excitatory projection neurons with distinct projection targets (Arszovszki et al., 2014; Gergues et al., 2020; Jin and Lee, 2021; Kim and Cho, 2017; LeGates et al., 2018; Wang et al., 2016; Wee and MacAskill, 2020; Xu et al., 2016), and a diverse population of inhibitory interneurons (Pelkey et al., 2017), including those expressing parvalbumin (PV), vasoactive intestinal peptide (VIP), and somatostatin (SST). These interneuron subtypes exhibit unique anatomical connectivity and exert differential control over specific circuit functions, including feedforward and feedback inhibition(Blasco-Ibáñez and Freund, 1995; Pouille and Scanziani, 2004), modulating spike-timing (Park and Kwag, 2012; Royer et al., 2012; Strüber et al., 2022), and establishing neuronal rhythms(Amilhon et al., 2015; Antonoudiou et al., 2020; Bartos et al., 2007; Klausberger et al., 2005; Losonczy et al., 2010; Stark et al., 2014). Interestingly, these interneurons are also differentially connected to projection-specific vCA1 outputs (Lee et al., 2014; Lodge et al., 2023). Although the role of VIP interneurons in vCA1 anxiety-related computations remains unclear, recent work implicates vCA1 PV interneurons in encoding anxiogenic states (Forro et al., 2022; Li et al., 2024; Tiwari et al., 2024; Volitaki et al., 2024) and social memories (Deng et al., 2019), while SST interneurons exhibit beta-band synchrony with amygdala SST cells during exploration of anxiogenic environments (Jackson et al., 2024) and can modulate theta oscillations during exploration (Mikulovic et al., 2018). Additionally, SST interneuron activity increases during extinction in both reward and fear conditioning paradigms, and their manipulation alters behavioral responses to conditioned contexts after extinction (Lacagnina et al., 2024; Li et al., 2024). However, how these cell types may differentially encode environmental features, and behavioral or emotional states to shape vCA1 output during the exploration of threatening environments is still being uncovered.

In the present study, we aimed to elucidate how distinct subtypes of vCA1 interneurons represent features of anxiety-related behavior, specifically examining their roles in differentiating between exploratory approach and avoidance decisions. We hypothesized that distinct features of anxiety-provoking experiences, such as features of the environment, behavioral states of avoidance, or internal emotional states, may be differentially represented in PV, VIP, and SST cells. Our findings highlight a striking functional specialization among interneuron classes within vCA1. Specifically, we found that when compared with PV and VIP neurons, vCA1 SST interneurons were specifically recruited during the execution of avoidance behaviors and were uniquely capable of generating predictive signals regarding future behavioral decisions related to approach vs avoidance. Remarkably, we observed that silencing vCA1 SST neurons impaired the efficiency of approach-avoidance decision-making, prolonging the time animals spent at the choice point before committing to a behavioral response. These results reveal a highly specialized role for SST interneurons in encoding behavioral states and decision variables linked to anxiety.

## Results

### Distinct recruitment of vCA1 interneuron subtypes during approach and avoidance

To examine how vCA1 interneuron subtypes are engaged during the exploration of anxiety-provoking environments, we expressed GCaMP8m in PV-, VIP- or SST-Cre mice and used fiber photometry to record population calcium signals in the elevated plus maze (EPM) (Fig 1A). We grouped trajectories into two types: approach trajectories, where the mouse travels from the end of one closed arm to the entrance to the center of the maze and then makes the decision to explore one of the open arms, and avoidance trajectories, where the mouse travels from the closed arm to the entrance to the center compartment and then makes the decision to escape to the opposite closed arm. Recordings revealed differential dynamics across interneuron populations across these two trajectory types. Both PV and VIP interneurons were robustly activated as animals entered the open arms and were less active than SST neurons during the retreat to the closed arms (Fig. 1B), indicating preferential recruitment by active exploration of anxiogenic regions, and consistent with previous reports (Forro et al., 2022; Li et al., 2024; Volitaki et al., 2024). Surprisingly, in contrast, SST-interneurons showed the exact opposite pattern: these neurons were selectively engaged during avoidance, exhibiting significant increases in population activity when the mouse retreated to the closed arms, but remained low during open-arm exploration (Fig 1B). Thus, SST interneurons encoded approach and avoidance states in a manner that was distinct from PV and VIP interneurons.

**Figure 1.**
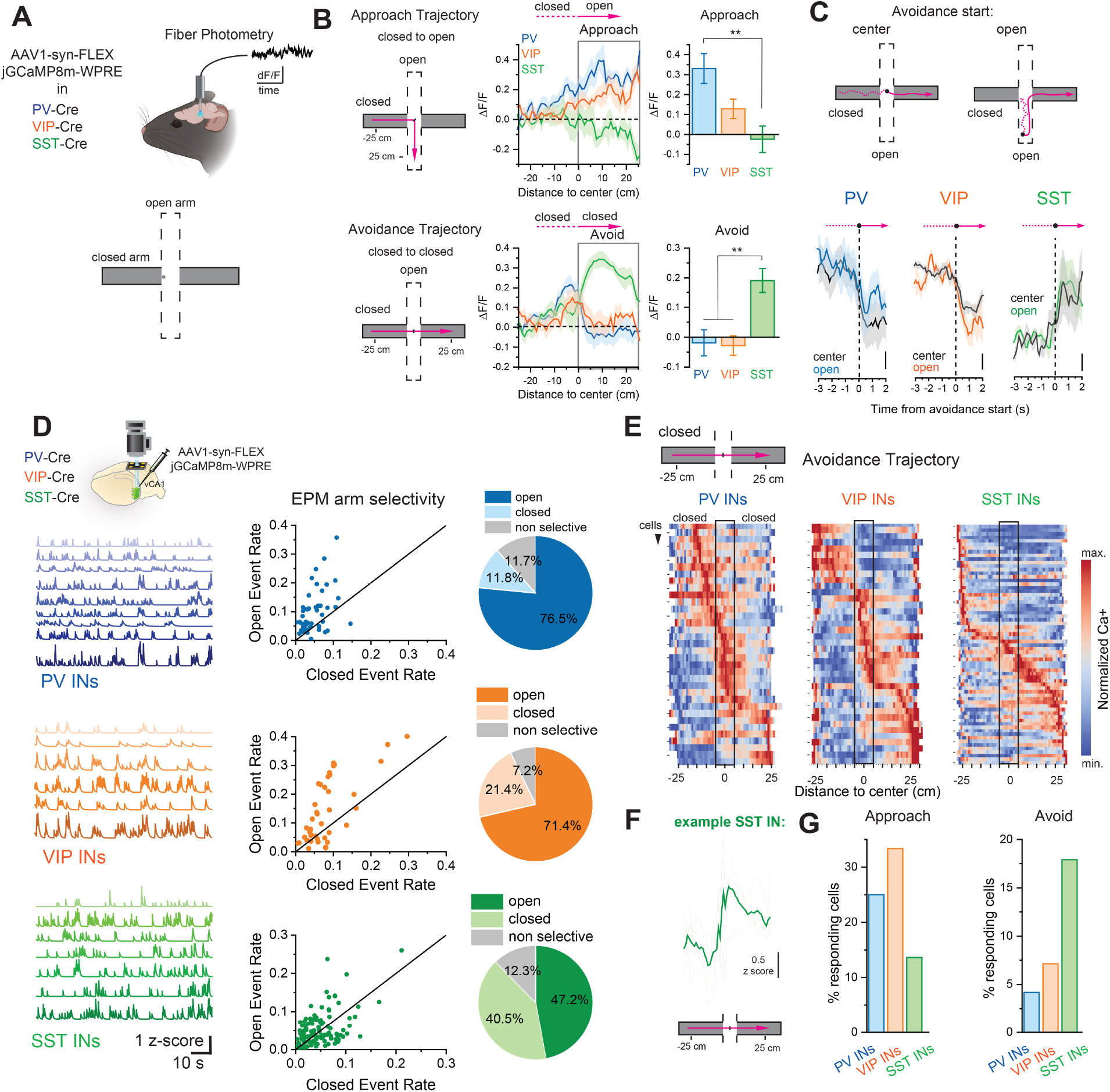
vCA1 somatostatin interneurons show a selective increase in activity during avoidance behaviors in the elevated plus maze. **A**. Schematic of photometry experiments in PV-, VIP-, and SST-Cre mice to allow for cell-type specific bulk population calcium recordings during the elevated plus maze (EPM). Calcium recordings done using viral expression of AAV1-syn-FLEX-GCaMP8m. **B**. Spatially-binned calcium traces (1 cm bins) across either approach trajectories or avoidance trajectories averaged across mice (PV: n=7; VIP: n=7; SST: n=9). Average activity values during the open-arm approach (0-25 cm, approach trajectory) showed an increase in PV and VIP and no noticeable activity in SST interneurons (one-way ANOVA, F = 7.47, p = 0.0046, Bonferroni-corrected post hoc PV vs VIP: t = -2.27, p = 0.1084, VIP vs SST: t = -1.65, p = 0.3515, PV vs SST t = -5.99, p = 0.0039). Conversely, average activity values during the avoidance into the opposite closed arm (0-25 cm, avoidance trajectory) show an increase in SST interneuron activity exclusively ((one-way ANOVA, F = 10.25, p = 8.58 × 10^-4^, Bonferroni-corrected post hoc PV vs VIP: t = -0.15, p = 1, PV vs SST: t = 3.75, p = 0.0037, VIP vs SST: t = 3.92, p = 0.0025). **C**. Comparing the time-locked response of avoidance behavior originating in the center portion of the maze to those originating at least 10 cm out in the open arm shows a similar temporal response at the onset of these behaviors across the interneuron classes. **D**. Example calcium activity traces of interneuron-specific GCaMP8m recordings in PV, VIP, and SST interneurons. Comparing event rates in the closed and open arms of the maze within single cells shows that the PV and VIP populations are enriched in cells selective to the open arm of the maze, while SST interneurons show a more heterogeneous response profile, but a large portion are selective to the closed arm of the maze (PV: 51 cells, 6 mice = 76.5% open arm responsive, 11.8% closed arm responsive; VIP: 42 cells, 4 mice = 71.4 % open arm responsive 21.4% closed arm responsive SST: 111 cells, 7 mice = 40.5% closed arm responsive, 47.2% open arm responsive. Pairwise chi-square tests, PV vs VIP: χ^2^(2) = 1.92, p = 0.38; PV vs SST: χ^2^(2) = 14.67, p = 6.5 × 10^-4; VIP vs SST: χ^2^(2) = 7.41, p = 0.025). **E**. Spatially-binned (1 cm) single cell averaged calcium responses to avoidance trajectories showing a large population of PV and VIP interneurons increasing their response towards the approach to the center of the maze with a large population of SST interneurons showing an increased response in and during the avoidance of the center towards the opposite closed arm. **F**. Example spatially-binned SST cell activity (individual trajectories, light green; averaged trace, dark green) during avoidance trajectories. **G**. Percentage of cells responding during either approach or avoidance trajectories. (Approach: PV = 13/51 cells; VIP = 14/42; SST = 16/111, Avoid: PV = 2/52; VIP = 3/42; SST = 20/111; chi-square test of independence across cell types, χ^2^(4) = 13.16, p = 0.010; pairwise 2 × 3 chi-square tests: PV vs VIP, χ^2^(2) = 1.51, p = 0.47; PV vs SST, χ^2^(2) = 7.57, p = 0.023; VIP vs SST, χ^2^(2) = 8.26, p = 0.016). ∗∗p < 0.01, ∗p < 0.05. All error bars indicate mean ± SEM.

To decouple approach and avoidance behavioral states from the sensory features of safe versus dangerous spatial cues, we compared the activity of these interneuron classes during two distinct types of avoidance behavior: 1) when the mouse retreated into the closed arm from the center, or 2) when the mouse retreated back to safety from the end of the open arm (Fig. 1C). If the dynamics were coding for the sensory features of the “safe” closed arm, we would expect distinct dynamics in the two trajectory types, as the safe closed arm is entered at different times (in scenario 2, the mouse still has to travel back along the open arm before reaching the closed arms). Within all interneuron subtypes, we observed identical dynamics regardless of the origin point of the avoidance-related retreat behavior: PV and VIP interneurons decreased their activity while SST interneurons showed a robust increase in activity at the onset of the avoidance behavior, regardless of where the avoidance was initiated.

To probe these dynamics in single neurons, we performed microendoscopic single-cell imaging of these interneuron classes (Fig. 1D). While we found some heterogeneity in the response profiles within each interneuron class, PV and VIP populations were overwhelmingly enriched in neurons selective for the open arm of the maze, while the SST population showed more closed arm-responsive neurons (Fig. 1D). Qualitatively examining single-cell responses across avoidance trajectories (Fig. 1E), we found that large populations of PV and VIP interneurons increased their activity as the mouse approached the center of the maze, whereas more SST interneurons increased their activity in the center and during retreat toward the opposite closed arm. To further probe this difference, we quantified the selectivity of SST neurons during approach and avoidance trajectories. Compared with PV and VIP neurons, a larger fraction of SST neurons responded during avoidance trajectories and a smaller fraction responded during approach trajectories (Fig. 1F–G). Taken together, these results highlight a pronounced functional dissociation among vCA1 interneuron subtypes, with SST interneurons selectively tuned to avoidance states and PV/VIP neurons preferentially linked to the exploratory approach.

### vCA1 interneurons more reliably encode behavioral intent than spatial position

Dorsal CA1 place cells exhibit direction selectivity on linear tracks, firing only when the animal traverses a field in one direction (McHugh et al., 1996; Ziv et al., 2013). In the EPM’s closed arms, however, direction maps onto behavioral state, approach toward the anxiogenic center versus retreat to safety, implying that direction selectivity there may report approach– avoidance state rather than simple heading (Fig. 2). To test whether this selectivity exists in vCA1, we examined the ability of each vCA1 interneuron subtype to represent the animal’s position on EPM during approach/avoidance runs. Using support vector machine (SVM) classifiers (Stefanini et al., 2020), we decoded the spatial location using the population activity recorded from PV, VIP, or SST interneurons. We constructed two distinct classifiers: a spatial decoder that estimated the animal’s true position at each point in each run, and a trajectory decoder, one that takes into account direction selectivity, and thus approach/avoidance state (Fig. 2A). At the same physical spot, heading flips meaning (e.g., +10 cm can be approach on left→right runs but avoidance on right→left), so in the trajectory decoder we aligned opposite directions to a common start/end to map each bin to a single intent. Comparing the two dissociates what’s encoded: if ensembles primarily encode approach/avoidance states, the trajectory decoder should outperform the spatial one; if they encode location, the spatial decoder should win because approach/avoidance-alignment collapses distinct ends into the same bin (examples in Fig. S1).

**Figure 2.**
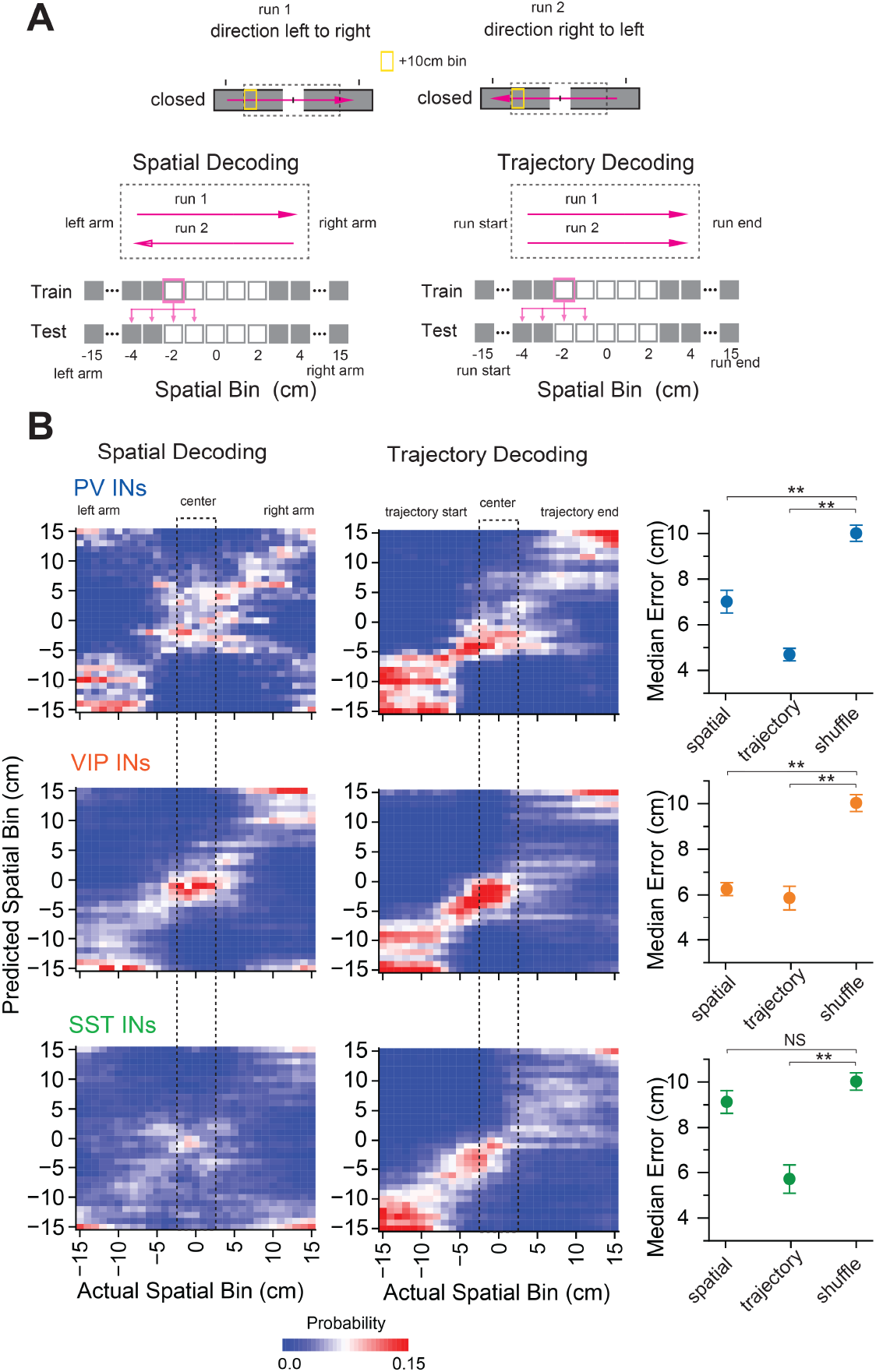
vCA1 SST interneurons more accurately track position along a behavioral trajectory than true space in the elevated plus maze. **A**. Schematic representation of trajectory decoding (left) and spatial decoding (right). For both decoders, a linear support vector machine classifier (SVM) using spatially-binned neural activity was trained to predict the location of the mouse (see text) **B**. Heatmaps showing the probability of the decoder in its spatial prediction compared to the actual position of the mouse for either the trajectory decoder (left) or the spatial decoder (right). For a decoder trained on PV and SST interneurons, the trajectory decoder was significantly more accurate than the spatial decoder, while the decoder trained using VIP interneurons showed an equal spatial accuracy across both. Only in SST interneurons was spatial decoder at chance levels. (n=28 cells per group, see methods, PV average median spatial error (cm): trajectory = 4.69±0.28, spatial = 7.01±0.49, Shuffle = 10.01±0.35; VIP average median spatial error (cm): trajectory = 5.86±0.52, spatial = 6.25±0.28, Shuffle = 10.02±0.37; SST average median spatial error (cm): trajectory = 5.71±0.62, spatial = 9.12±0.49, Shuffle = 10.02±0.38. PV cells: Tukey-corrected post hoc test TD vs shuffle: q = 12.68, p < 10^-4, SpD vs shuffle: q = 7.48, p = 2.44 × 10^-6, VIP cells: Tukey-corrected post hoc TD vs shuffle: q = 9.74, p < 0.0001, SpD vs shuffle: q = 9.38, p < 0.0001, SST cells: Tukey-corrected post hoc, TD vs shuffle: q = 10.17, p < 0.0001, SpD vs shuffle: q = 2.03, p = 0.3239)) ∗∗p < 0.01, ∗p < 0.05. error bars indicate mean ± SEM.

Notably, only in SST interneurons did the spatial decoder fail to perform better than chance, whereas the trajectory decoder remained significantly above chance (Fig. 2B). This pattern of direction-dependent selectivity indicates that SST activity tracks behavioral intent tied to heading within the closed arms, approach toward the anxiogenic center versus retreat to safety, rather than absolute spatial position. By contrast, in VIP and PV interneurons both spatial and trajectory decoders performed significantly above chance, consistent with a more mixed code that carries both positional and intent-related information rather than selectively emphasizing approach-avoidance state.

### SST interneurons prospectively signal impending avoidance decisions

As our data indicated that SST neurons preferentially encode avoidance states, we next asked when these signals arise and whether they predict behavioral choices. Specifically, we tested whether vCA1 interneuron populations carry prospective signals about the animal’s upcoming decision to enter the open arms or return to the closed arms. To investigate this, we analyzed changes in neural activity, measured with fiber photometry, during trajectories as the mouse moved toward the center of the maze. For each trial, we computed the change in activity from the beginning of the trajectory to the center-entry point immediately before the animal made its avoidance/exploration choice (Fig. 3A). SST interneurons showed a greater ramp in activity on trajectories that ended in avoidance compared with those that ended in open-arm exploration (Fig. 3B). In contrast, changes in PV and VIP activity did not differ based on the mouse’s subsequent decision. These findings raise the possibility that vCA1 SST interneurons compute a prospective signal that biases subsequent avoidance choices.

**Figure 3.**
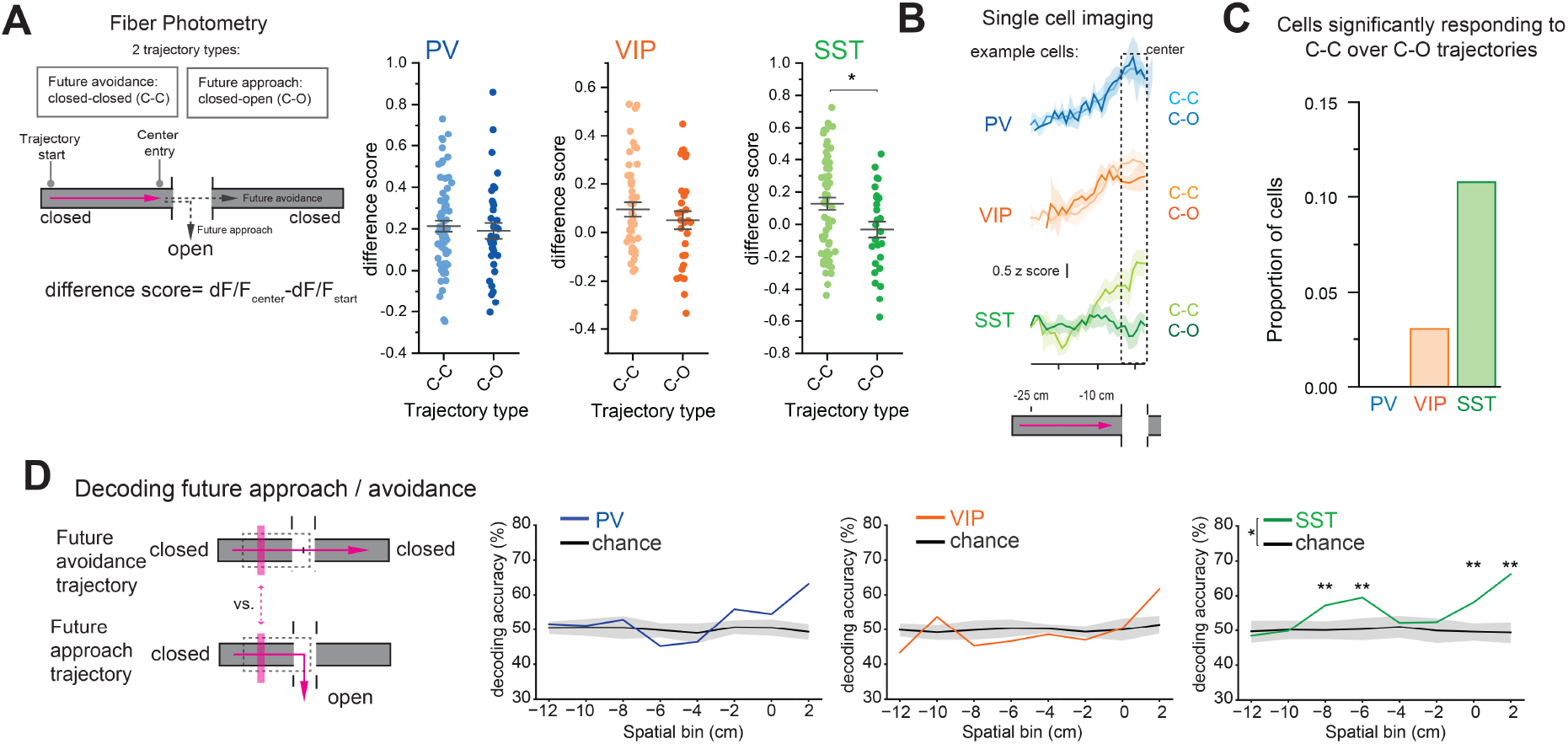
SST interneurons uniquely exhibit a predictive signal of future avoidance behaviors. **A**. Schematic for analysis of change in activity from start to choice point in closed-closed vs closed-open trajectories **B**. Fiber photometry recordings across all trajectories found that while PV and VIP interneurons showed no difference in the ramping of their activity across these trajectory types, SST interneurons showed an increase in activity prior to avoidance trajectories. (n=7 PV mice, 7 VIP mice and 9 SST mice, unpaired t-test PV: CC vs CO, t = 0.49, p = 0.62; VIP: CC vs CO, t = 0.92, p = 0.36; SST: CC vs CO, t = 2.46, p = 0.016.) **C**. Single cell imaging data identified a higher proportion of SST interneurons showed ramping as the mouse reached the center of the maze prior to future avoidance trajectories compared to future approach trajectories. (PV 0/48 cells, VIP 1/42, SST 8/75; chi-square, χ^2^(2) = 7.49, p = 0.0236) **D**. Schematic of trajectories leading to future approach versus avoidance choices as the mouse reaches the center of the maze, used for classifier training. While decoders trained on PV and VIP interneurons could only distinguish activity after the behavioral approach/avoid decision had been made, SST interneurons could predict the future trajectory type prior to the behavioral decision. Because only SST showed a significant overall effect in a one-way ANOVA comparing true vs shuffle decoding across spatial bins (PV: F = 1.50, p = 0.24; VIP: F = 0.05, p = 0.83; SST: F = 6.89, p = 0.02), we restricted bin-wise tests to SST. Post hoc t-tests comparing SST vs shuffle at each spatial bin: −12: t = −0.61, p = 0.5500; −10: t = −0.17, p = 0.8643; −8: t = 4.13, p = 0.0014; −6: t = 4.12, p = 0.0016; −4: t = 0.50, p = 0.6281; −2: t = 1.29, p = 0.2194; 0: t = 3.75, p = 0.0041; 2: t = 6.64, p = 5.8 × 10^-5^. ∗∗p < 0.01, ∗p < 0.05. All error bars indicate mean ± SEM. Shading in chance lines indicates ± SD.

Single-neuron recordings corroborated these results. As mice approached the center decision point, we identified neurons whose ramping activity distinguished avoidance (closed→center→closed; C–C) from approach (closed→center→open; C–O) outcomes (Fig. 3B). Quantitatively, when compared to PV and VIP, SST interneurons contained the highest fraction of cells with significantly stronger responses on C–C than C–O runs, further underscoring their distinctive role in establishing vCA1 network states that bias impending choices toward avoidance.

To probe these dynamics at the population level, we trained SVM classifiers using calcium activity from ensembles of each interneuron subtype during trajectories that ended in either future approaches or avoidance. We then determined where along the trajectory the neural activity could reliably distinguish between upcoming behavioral choices (Fig. 3D). Consistent with their non-predictive activity profiles, decoders based on PV and VIP interneuron populations were unable to distinguish trajectory type until after the behavioral choice was made. In contrast, the decoder based on SST interneuron calcium activity successfully predicted future approach versus avoidance decisions several centimeters before the center decision point. Collectively, these results indicate that SST interneurons uniquely carry anticipatory information about avoidance, reflecting the animal’s behavioral intent within the vCA1 network before a behavioral commitment is executed.

### SST inhibition prolongs threat-related decisions

We next asked how SST activity shapes approach–avoidance behavior. To test this causally, we examined whether inhibiting SST neurons influences the execution of approach-avoidance decisions in the EPM. We bilaterally expressed the optogenetic silencer stGtACR2 in vCA1 SST interneurons and transiently inhibited them as mice entered the decision point at the center of the EPM (Fig. 4A). This manipulation significantly extended decision time, increasing the latency to choose between approach and avoidance (Fig. 4B). Under SST inhibition, mice spent more time in the center, consistent with prolonged conflict at the choice point. These results suggest that SST interneurons facilitate rapid avoidance decisions, and that their disruption delays conflict resolution at the decision point.

**Figure 4.**
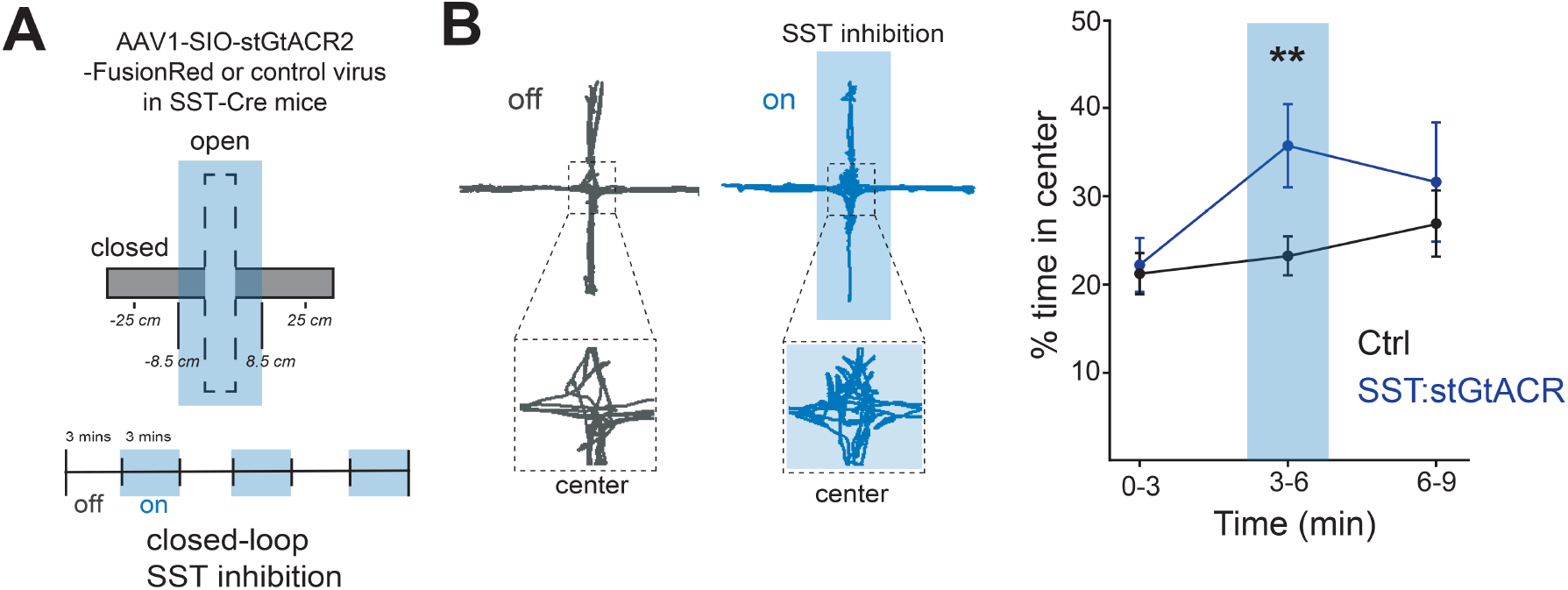
Inhibition of vCA1 SST interneurons prolongs threat-related decisions. **A**. Schematic of bilateral vCA1 SST-stGtACR2 center/open-triggered photoinhibition upon entry into the decision point of the EPM. **B**. Time spent in the center across 3-min epochs during light-off versus light-on periods. Right, example trajectory from an SST-stGtACR2 mouse showing prolonged center occupancy during the light-on epoch with repeated nosepokes into open and closed arms, consistent with extended decision making. SST-stGtACR2 mice spent more time in the center during the 3–6 min epoch compared with mCherry controls, consistent with prolonged decision latency (two-sample t-tests, SST-stGtACR2 n = 6 vs mCherry n = 9; 0–3 min: t = 0.26, p = 0.7983; 3–6 min: t = 2.67, p = 0.0189; 6–9 min: t = 0.66, p = 0.5207).

## Discussion

This work identifies a cell-type-specific mechanism in vCA1 by which SST-expressing interneurons represent internal avoidance-related variables that can prospectively guide anxiety-related behavioral choices. It builds on prior work showing that vCA1 encodes discrete stimulus identities (Biane et al., 2025, 2023) and emotional states (Xia et al., 2025), by demonstrating that vCA1 population activity also tracks moment-by-moment avoidance states. Using a combination of optical tools, we show that SST interneurons: **1)** are selectively recruited during avoidance behavior, in contrast to PV and VIP interneurons, which are activated during exploratory approach; **2)** ramp their activity in anticipation of avoidance decisions; **3)** encode a prospective signal that predicts future avoidance behavior; and **4)** are required for efficient approach-avoidance decision-making and risk assessment. These findings position SST interneurons as a critical node for computing internal states that bias decisions under anxiety.

The distinct patterns of recruitment among vCA1 interneuron subtypes reveal complementary roles in encoding internal states and behavioral strategies during the exploration of anxiogenic environments. PV and VIP interneurons were preferentially active during approach behaviors, with PV responses likely reflecting feedforward activation from local pyramidal neurons, a known property of these neurons in CA1 (Ali et al., 1998; Bland et al., 1988; Huh et al., 2016; Klausberger and Somogyi, 2008; Strüber et al., 2022; Sun et al., 2014; Ylinen et al., 1995). These interneurons may act to constrain excitability during exploration and approach drives, stabilizing network output and potentially limiting maladaptive overactivation in threatening contexts, as has been recently shown (Volitaki et al., 2024). In contrast, during avoidance and retreat from threatening spaces, PV and VIP activity diminishes sharply and SST activity ramps up, a switch that may facilitate transitions between competing behavioral states. Given known reciprocal inhibition between PV and SST interneurons in CA1 (Brzdąk et al., 2023), SST recruitment may actively suppress approach-related network activity and promote an avoidance-related behavioral state.

A key finding of this study is that SST interneurons gate competing approach and avoidance signals, shaping behavior at conflict-related decision points. As animals approach the maze center, where risk assessment occurs, SST activity ramps, but notably only in trials that ultimately result in avoidance. Silencing SST interneurons during this period of risk assessment abolishes the ability of the vCA1 population to prospectively represent future behavioral outcomes. This supports a role for these interneurons in modulating risk assessment states in vCA1 output neurons, sculpting an output signal that promotes avoidance over approach.

This network mechanism, which balances approach and avoidance during risk assessment, is particularly relevant to the neural basis of anxiety, especially heightened threat sensitivity and behavioral inhibition (Jovanovic and Ressler, 2010; Kheirbek et al., 2012). The ramping activity of SST interneurons, seen only on trials that end in avoidance, indicates that they encode internal threat signals that bias future choices, a dynamic consistent with predictive-coding models in which internal threat estimates guide behavior to minimize aversive outcomes. The predictive signal in SST interneurons likely arises from the convergence of contextual and emotionally salient inputs. Potential candidate inputs could arise from intrahippocampal pathways and projections from the amygdala, midline thalamus, or medial septum (Ali et al., 1998; Felix-Ortiz et al., 2013; Haam et al., 2018; Honoré et al., 2021; Jackson et al., 2024; Maccaferri, 2005; Sun et al., 2014), enabling these cells to sculpt vCA1 activity during threat evaluation. Supporting this, recent work has shown increased coherence between SST interneurons in vCA1 and the amygdala during avoidance (Jackson et al., 2024), implicating them in a distributed circuit coordinating threat-related states. These findings suggest that SST interneurons act not only as local modulators of hippocampal dynamics but also as integrators of predictive state information from extended limbic networks; dysregulation of which could drive maladaptive avoidance and excessive anticipatory anxiety (Abdallah et al., 2017; Kirkby et al., 2018; Lazarov et al., 2017).

Collectively, these results reveal a microcircuit in which vCA1 SST interneurons encode internal avoidance-related signals and influence risk assessment under anxiety. By demonstrating cell-type–specific representations of internal behavioral drives, our findings broaden traditional models of hippocampal function and highlight SST interneurons as a promising target for modulating maladaptive avoidance in anxiety disorders.

**Figure S1.**
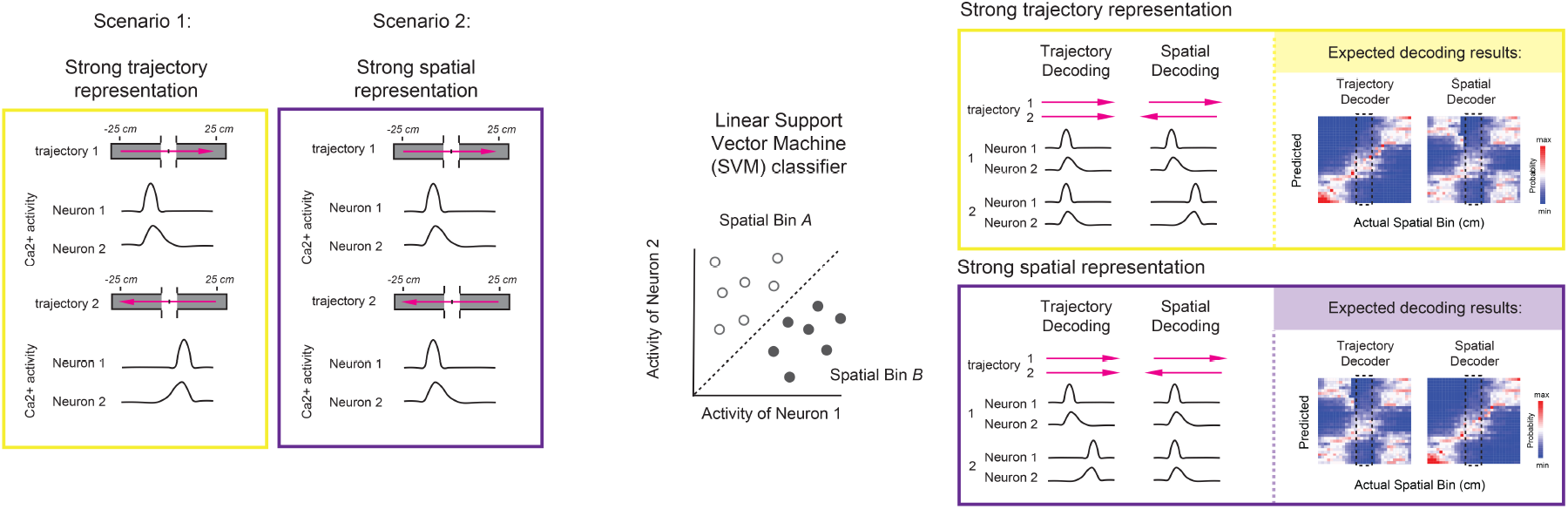
Schematic of trajectory and spatial decoder approaches.

## Methods

### Animals

All animal and experimental procedures were approved by the Institutional Animal Care and Use Committee (IACUC) at the University of California, San Francisco and carried out in accordance with the National Institutes of Health *Guide for the Care and Use of Laboratory Animals*. The following transgenic mice were obtained from The Jackson Laboratory and used in this study: PV-Cre (JAX strain #: 017320), VIP-IRES-Cre (010908), and SST-IRES-Cre (013044). Homozygous mice of each of these strains were crossed with wild type C57BL/6 mice from The Jackson Laboratory such that all experimental mice were heterozygous for Cre recombinase. Adult mice were used for all experiments, ranging between 3-8 months old. Mice were group housed (between 2-5 littermates) in a temperature (22-24°C) and humidity (40-60%) controlled environment and kept on a 12 hour light/dark cycle. All experiments were carried out during the light cycle.

### Surgical Procedures

Surgical procedures were carried out on adult mice aged between 12-20 weeks. In a head-fixed stereotaxic frame (David Kopf), mice were anesthetized with 1.5% isoflurane supplied with an oxygen flow rate of 1 L min−1. Eyes were lubricated with an ophthalmic ointment, and body temperature was maintained at 35-37 °C with a warm water recirculator (Stryker). Around the top of the head, the fur was shaved to expose the skin which was then sterilized by alternating between three rounds of isopropyl alcohol and betadine solution application. At the incision site, lidocaine HCl was injected subcutaneously. Meloxicam and slow-release buprenorphine were provided for analgesia and all mice were closely monitored for post-surgical care.

For viral injection and optic fiber / lens implants, the skull was exposed, scored, and craniotomy then carried out using a 0.5 - 1.0mm drill bit (David Kopf). For all calcium imaging experiments, GCaMP8m was used. For interneuron specific recordings, AAV1-syn-FLEX-jGCaMP8m-WPRE (titer: 2.4 × 10^13; Addgene, catalog number: 162378) was used and diluted 1:3 in sterile 1X PBS.

For pan-neuronal imaging, AAV1-syn-jGCaMP8m-WPRE (titer: 2.4 × 10^13; Addgene, catalog number: 162375) was used and diluted 1:3 in sterile 1X PBS.. Both virus injections and fiber/lens implants were performed during the same surgery. First, virus delivery was performed using a Nanoject III syringe (Drummond Scientific) with a fine tip (width 20-30 µm) glass needle at a rate of 10 nanoliters/second. A volume of 167 nL of virus was delivered to the following coordinates (in mm from bregma) for vCA1 targeting: -3.16 A/P, -3.20 M/L, at depths from brain surface -3.75, -3.50, and -3.25 D/V. Between each D/V injection site the virus was allowed at least 5 minutes to settle before slowly moving the glass needle to the next injection site. For bilateral optogenetic silencing experiments, AAV1-hSyn1-SIO-stGtACR2-FusionRed (Addgene, 105677) or a control virus (AAV1-hsyn-mCherry (Addgene, 114472) was injected bilaterally at 167.7 nL per D/V site. For photometry experiments, a 5.5mm in length 400 µm borosilicate optic fiber (Doric Lenses, numerical aperture 0.66) was used. For microendoscopy experiments, a GRIN lens (either 0.5, 0.6, or 1.0mm diameter, Inscopix) was used. The majority of the GRIN lenses used had an integrated baseplate (Inscopix), while in some animals a baseplate was manually attached using further dental cement at least three weeks later. In animals without an integrated baseplate, the exposed GRIN lens was protected by sealing a protective rubber mold around the top of the lens such that mice could be socially housed without causing damage to the lens. All implants were done on the left hemisphere. For bilateral optogenetic silencing experiments, 200 µm optic fibers were created as previously published (Jimenez et al., 2018)and implanted bilaterally. All implants were fixed to the skull through the use of 1-2 skull screws and Metabond adhesive cement (Parkell). All animals were given at least 4 weeks after surgery before being used for behavioral and recording experiments.

## Behavior

### Elevated Plus Maze

On the day of recording, mice were transferred to the room where behavioral experiments were performed and allowed at least 60 minutes of habituation to the room. Mice were then placed in a standard EPM maze with the platform elevated 30.3 cm off the floor and the following maze dimensions: each full arm length of 60 cm, arm width of 5 cm, closed arm wall height of 17.8 cm, and open arm ledges of 1.2 cm. Room lighting was adjusted such that the open arms were between 400-500 lux and closed arms between 150-250 lux. Mice were first placed into the end of one of the closed arms and allowed 18 minutes to explore the maze while behavior was recorded with a webcam (Logitech; Lausanne, Switzerland). The maze was thoroughly cleaned with 70% ethanol solution and allowed to dry between different mice. In real time, mice behavior was tracked using Ethovision XT 10 software (Noldus, Leesburg, VA) and all tracking was manually inspected following recordings to ensure accuracy before moving onto further behavioral analysis. Personnel inspecting animal behavior were blinded to experimental groups.

### Closed-Loop Optogenetics

In the experiments utilizing behaviorally closed-loop optogenetic silencing of SST interneurons, the optogenetic areas of the maze were established using Ethovision XT 10 software. For the elevated plus maze, the optogenetic “on” area was set to cover the entire open arms of the maze and the last 8.5 cm of each of the closed arms near the center of the maze to ensure that the optogenetic silencing was being performed at a time that the mouse made its decision while getting towards the center of the maze. The animal was live tracked and in alternating 3 minute windows the optogenetic “on” rule was followed, allowing for within animal control of the effects of optogenetic silencing.

### Photometry Recordings

At least four weeks following virus injection and optic fiber implant, mice were subjected to behavior tasks during recording of photometry signals. Mice were habituated to having the optic fiber plugged in while freely moving in a home cage for at least one day prior to experimental trials. On the days of recording, the exposed optic fiber was plugged into a connector fiber optic cable and each mouse was allowed 10 minutes of habituation before being placed into the behavioral apparatus. Fluorescence signal was recorded using the Tucker-Davis Technologies fiber photometry system and Synapse software. The GCaMP signal was recorded using a 465 nm LED (Doric Lenses) and an isosbestic signal recorded using 405 nm LED (Doric Lenses). Recordings were performed at 210 Hz and 330 Hz, respectively. For all mice, the final output of the GCaMP fluorescence intensity was set between 150-250 mV. Photometry signals were processed using the Python toolbox GuPPy (Sherathiya et al., 2021), subtracting the scaled isosbestic signal from the GCaMP signal, and final data reported as dF/F.

### Freely moving Ca2+ imaging

At least four weeks following surgery mice were checked for GCaMP expression using the attachment of the nVoke miniaturized microscope (miniscope; Inscopix, Palo Alto, CA). The microscope was attached to the magnetic baseplate and secured in place with a small screw. The fields of view for each mouse were explored as they were freely moving in a new cage and up to three different z planes were chosen for final imaging of the mouse and used across all imaging days. On the day of experimental imaging, mice again had the minscope attached and allowed 10 minutes of freely moving habituation in their home cage. Calcium recording videos were obtained using the nVoke acquisition software (Inscopix) that was triggered by an external TTL pulse from the EthoVision XT 10 and Noldus box system to align behavioral and calcium videos. Videos were captured at a frame rate of 15 Hz spanning across three different z planes, with an individual plane being imaged at 5 Hz.

### Extracting Ca2+ signal

Calcium videos were preprocessed using Inscopix Data Processing Software (IDPS), including cropping of the videos to isolate the field of view, de-interleaving the separate z planes, spatially bandpass filtering, and a rigid motion correction. Additionally, some videos received further non-rigid motion correction based on template matching using NoRMCorre. Cell segmentation and calcium transient time series data were extracted using non-negative matrix factorization for microendoscopic data (CNMFe) which has been optimized for 1-photon GRIN lens Ca2+ imaging (Zhou et al., 2018). All putative neurons detected were carefully manually inspected for both appropriate spatial properties and Ca2+ dynamics. The resulting dF traces of individual cells were z scored to the entire duration of the recording. To detect Ca2+ events and inferred spiking rate, OASIS was used embedded within the CNMFe workflow.

## Data Analysis

### Trajectory detection

Trajectories were detected using both visual inspection of behavioral videos and the *x,y* coordinates of the center of the mouse’s body, as tracked by Ethovision XT 10 software. First, all behavioral videos were aligned such that the center of the maze was established as *x,y* = 0,0, with the closed arms extending along the x-axis and the open arms along the y-axis. To match the calcium data, all behavioral videos were downsampled to 5 Hz. To be considered a closed-closed trajectory, the mouse had to start its trajectory from the first half of a closed arm (at least 15 cm away from the center) and make it to the opposite side of the other closed arm (at least 15 cm away from the center). To be considered a closed-open trajectory, the mouse had to again start from the first half of the closed arm and now make it at least 6 cm into one of the open arms. This is a point where the back feet of the mouse have left the center of the maze. For detection of the start of avoidance behavior in the center or the open arm, the start time of a direct avoidance trajectory back into the closed arm (not stopping along the way) was manually determined.

### Trajectory Traces

Photometry/cell calcium data for each trajectory was spatially binned in either 1 or 2 cm bins and the average calcium signal in these spatial bins was used. Calcium data was interpolated along a trajectory for any spatial bins that might have been skipped due to the downsampling of the behavioral videos to 5 Hz. Time periods when mouse velocity was less than 2 cm/s for a period of 1 second were removed from the data before binning.

### Single cell selectivity

To compute single cell selectivity to the different compartments in the EPM, event rates were determined using all time bins when the animal was present in a given arm and moving above the threshold of 2 cm/s. The difference in these event rates was compared against a shuffled data set (1000 iterations) for each cell in which Ca2+ events were shuffled but the behavior time bins kept constant. A cell was designated as selective for a given arm if the event rate difference between the arms exceeded 1 SD from the shuffled distribution. Non-selective cells showed no significant difference between event rates and the shuffled distribution. Cells were designated as significantly responsive to approach or avoidance by comparing the average calcium signal in spatial bins during the approach epoch (−8 to 0 cm from the center) versus the avoidance epoch (0 to +8 cm), and classifying cells whose observed difference exceeded 1 SD of a shuffled distribution in which calcium events were permuted relative to behavior (Jimenez et al., 2018).

### Population level analysis

Linear support vector machine (SVM) classifiers were used to discriminate activity patterns of neural population data, as previously described (Biane et al., 2023). When multiple (2+) classes were used, multiple linear decoders were trained on pairs of discrete categories combined using majority-based error-correction codes. Pseudo-population recordings were generated by combining cell data across multiple mice within each group. For interneuron recordings, all pseudo-populations were balanced in cell number such that 28 cells were used across all groups and each iteration a random selection of available cells was chosen. To be included, cells had to see a minimum of 4 different individual trials within each class of trajectory. Trajectories were created by randomly selecting individual trajectories from within the pseudo-population of cells and balancing right-to-left and left-to-right trials. For decoding, a training-testing split of 50:50 was used. Cross-validation was used such that all trials were used for testing at least once. In total, we repeated pseudo-population generation and SVM analysis across 100 iterations of a given analysis. Final SVM decoding accuracy was reported as the average performance of the decoder across the 100 iterations.

## Resource availability

### Contact for reagents and resource sharing

Further information and requests for resources and reagents should be directed to and will be fulfilled by the Lead Contact, Mazen Kheirbek.

### Materials Availability

This study did not generate new unique reagents.

### Data and Code Availability

All analysis code is provided at https://github.com/mkheirbek. All the raw calcium imaging data will be provided upon request to the corresponding author.

## Acknowledgements

The authors would like to thank Vijay Namboodiri, Jeremy Biane and Alexandra Klein for discussion and comments, MAK was supported by NIMH (R01 MH108623, R01 MH117961, R01 MH136270, R01 MH125515), NIDCD (R01 DC019813) a One Mind Rising Star Award, a Research Grant from HFSP (Ref. No-RGY0072/2019), the Esther A. and Joseph Klingenstein Fund, the Pew Charitable Trusts, the McKnight Memory and Cognitive Disorders Award and The Ray and Dagmar Dolby Family Fund.

