## Supplementary figures and images for "vCA1 SST neurons represent avoidance states that guide anxiety-related behavioral choices"

### Figure S1

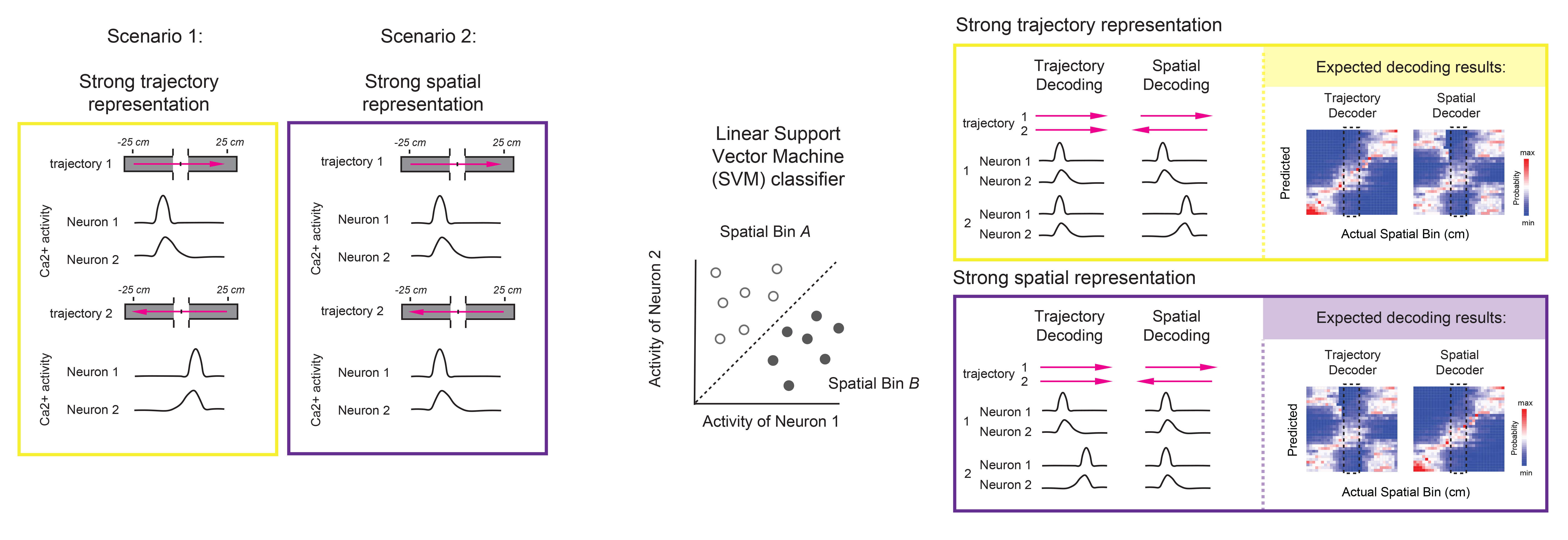
